## Supplementary figures and images for "In search of disentanglement in tandem mass spectrometry datasets"

### Figure_S1.png

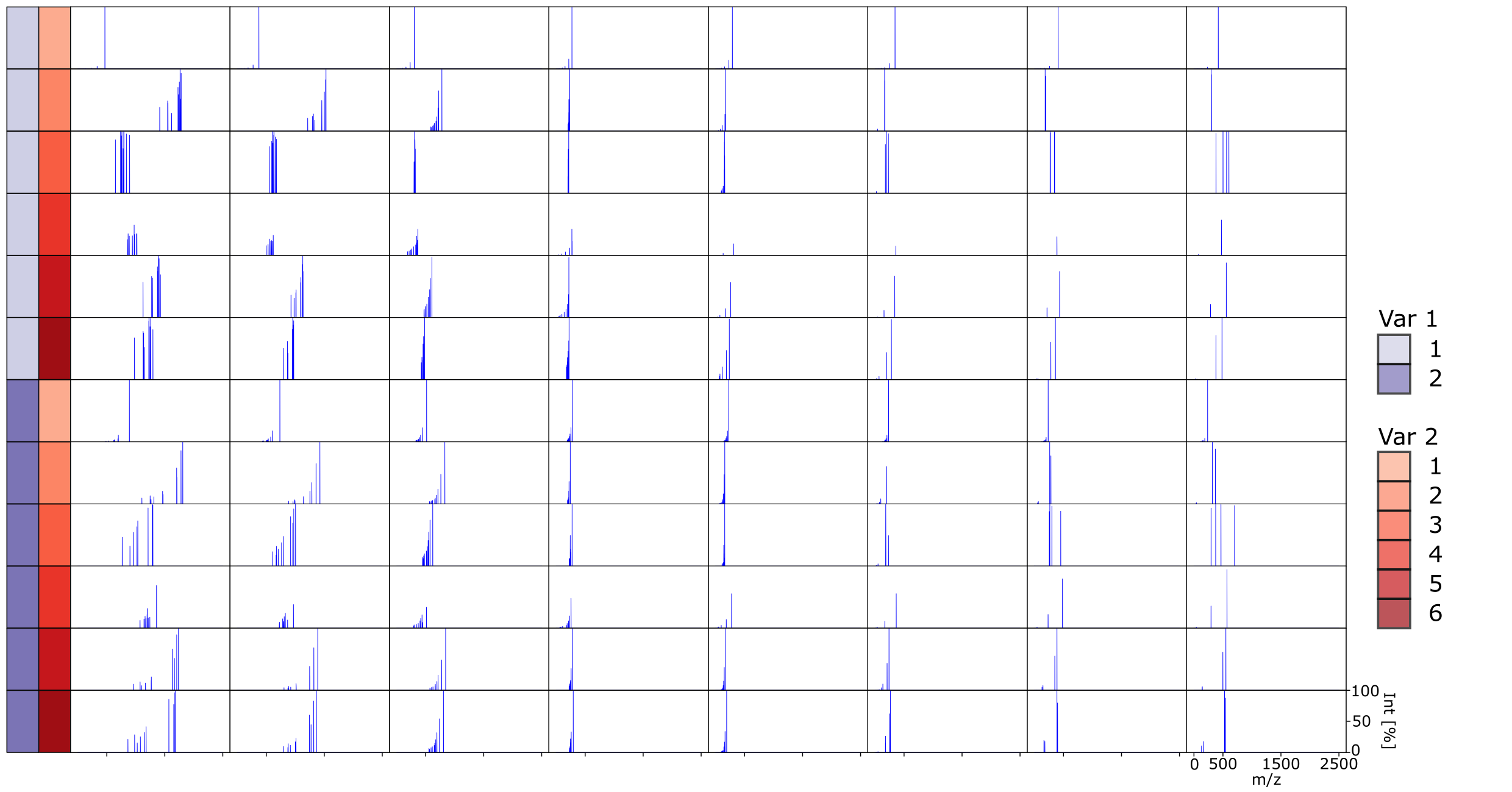

### Figure_S2.png

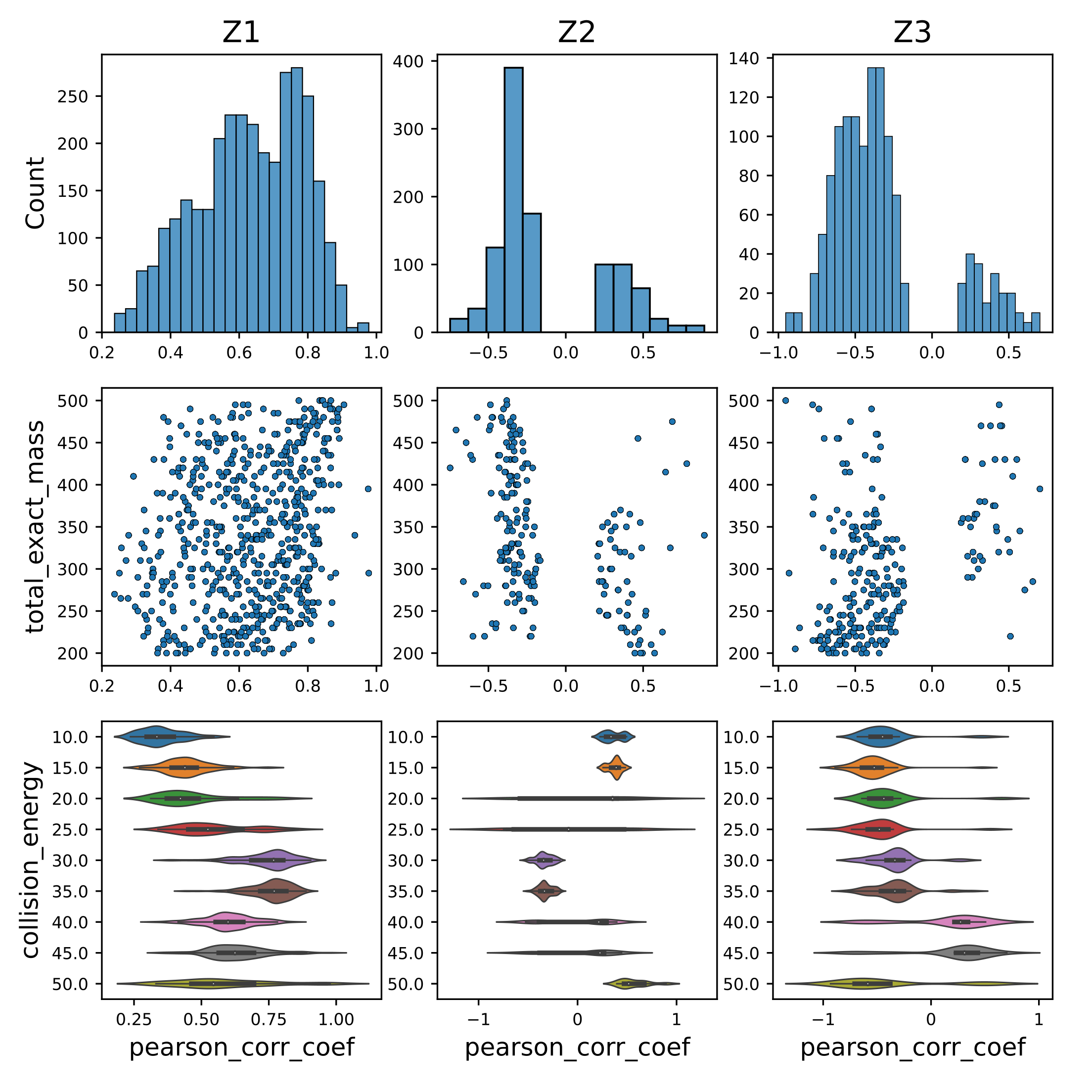

### Figure_S3.png

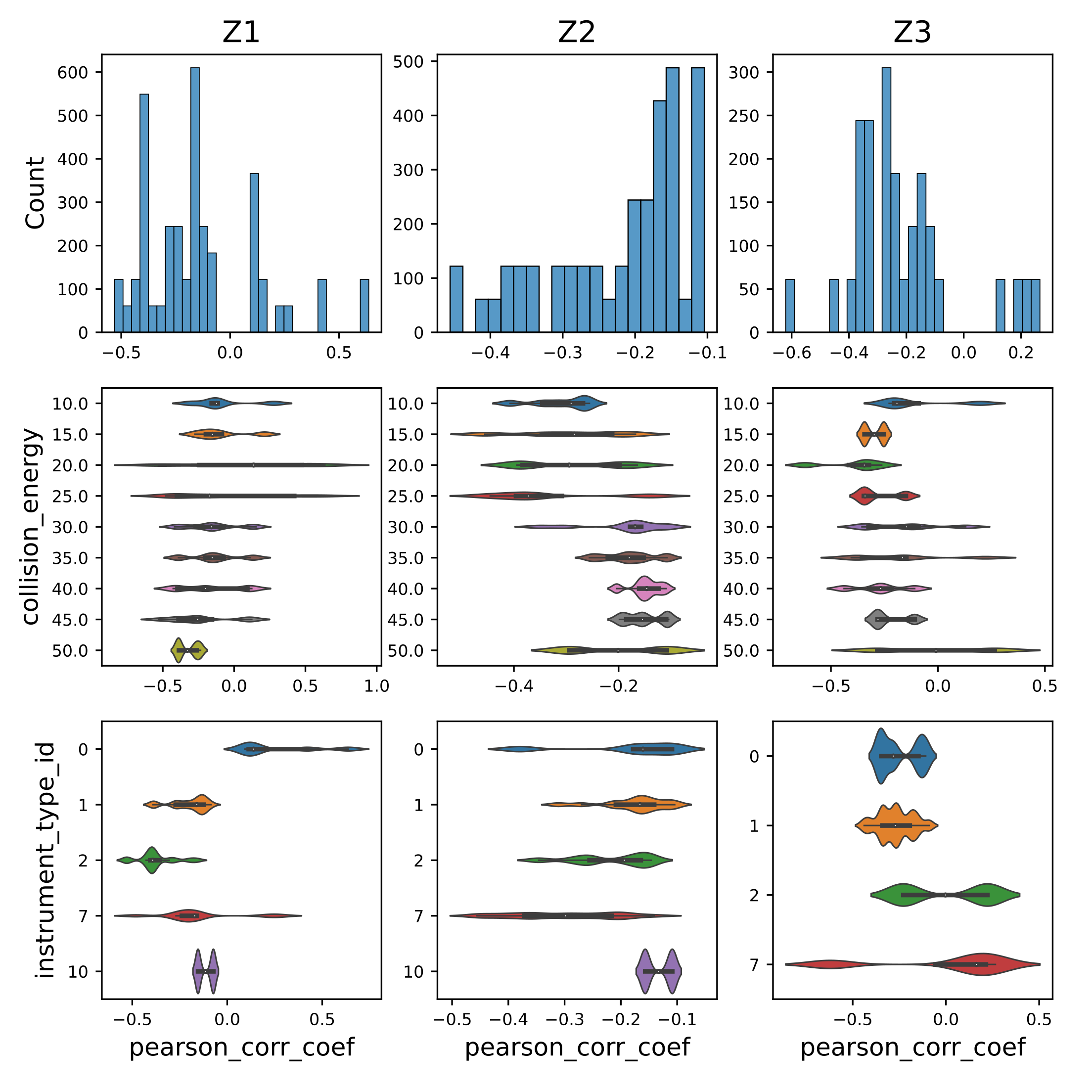

### Figure_S4.png

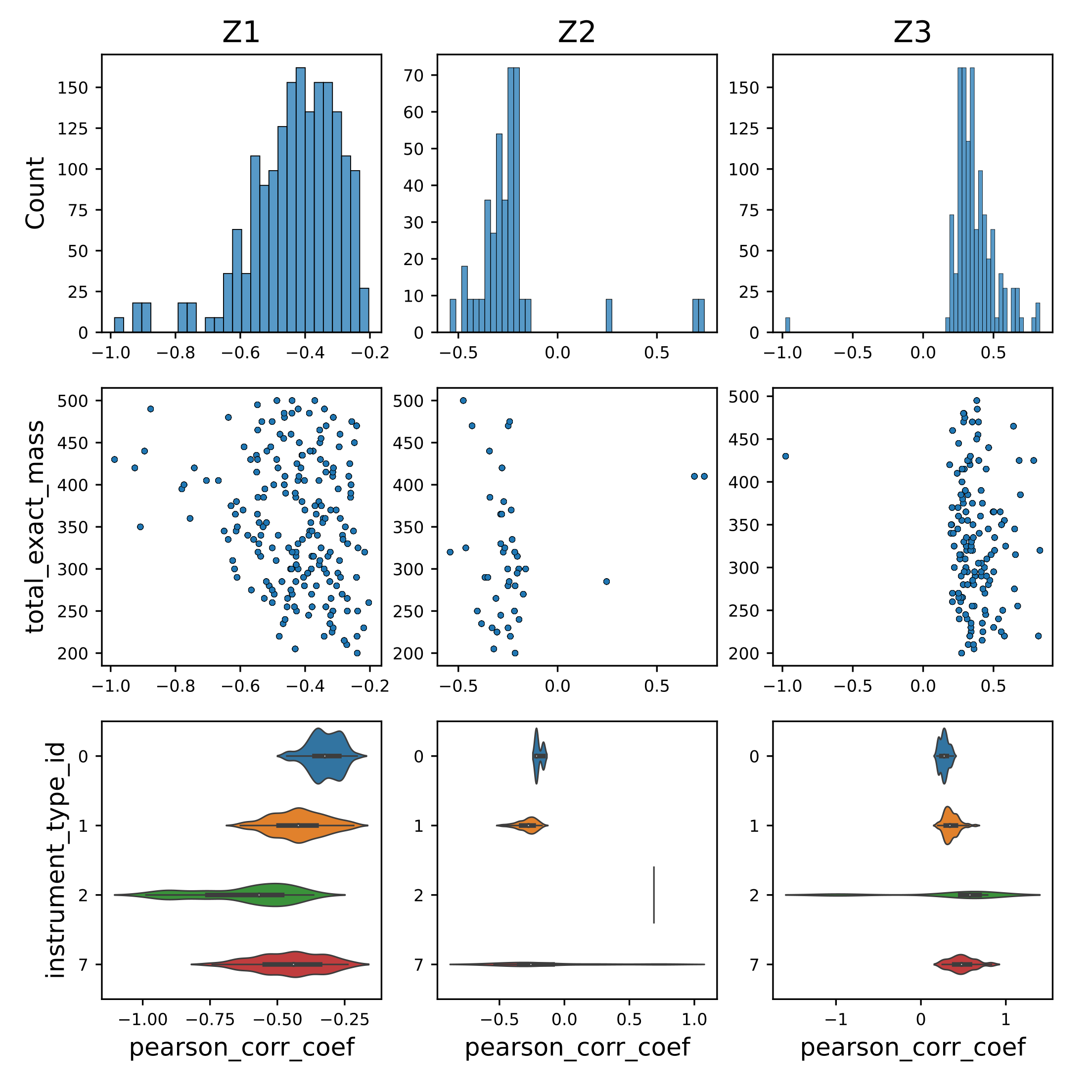

### Figure_S5.png

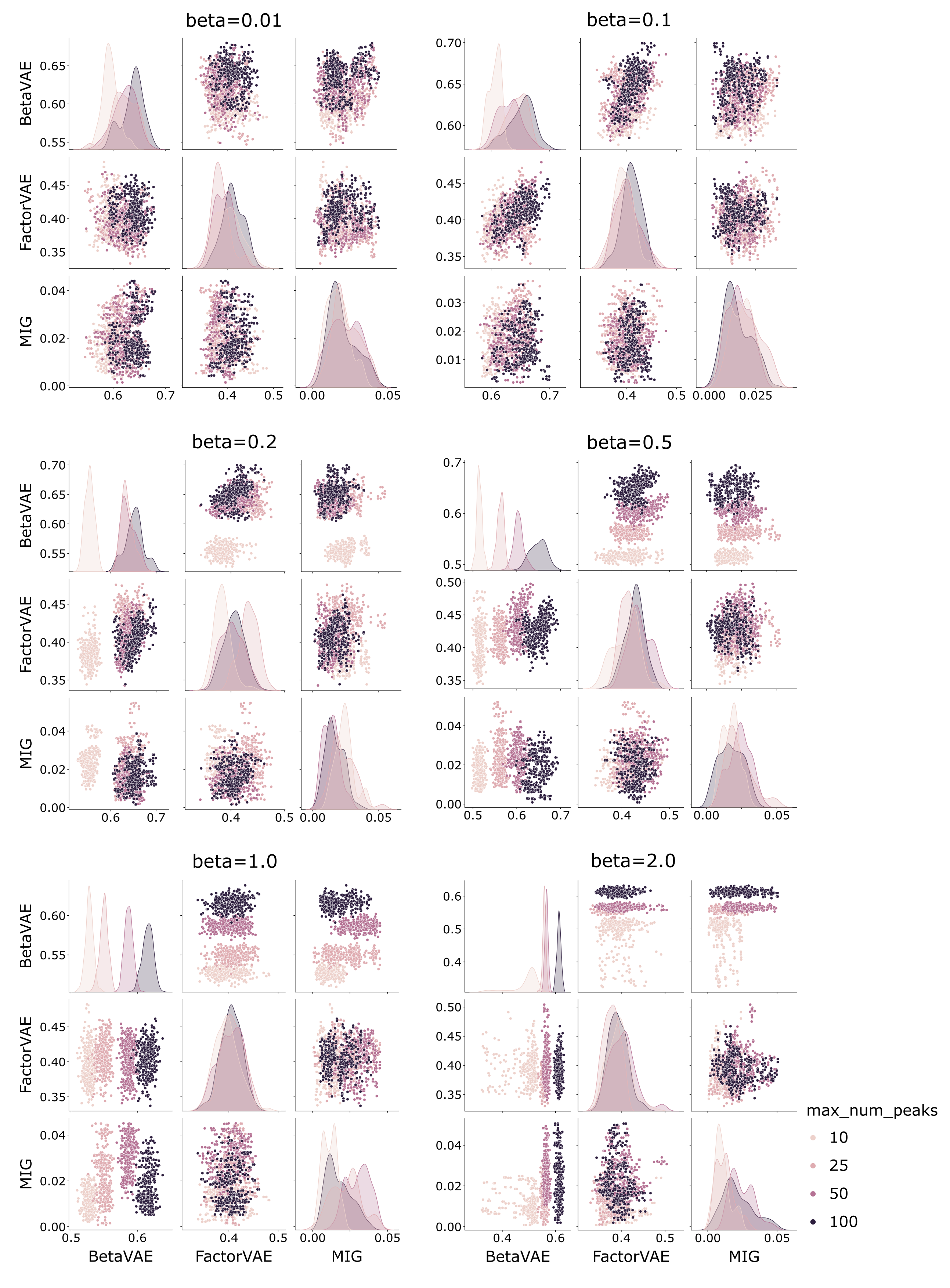
